# The respiratory cycle modulates distinct dynamics of affective and perceptual decision-making

**DOI:** 10.1101/2024.03.26.586076

**Authors:** Malthe Brændholt, Niia Nikolova, Melina Vejlø, Leah Banellis, Francesca Fardo, Daniel S. Kluger, Micah Allen

## Abstract

Respiratory rhythms play a critical role not only in homeostatic survival, but also in modulating other non-interoceptive perceptual and affective processes. Recent evidence from both human and rodent models indicates that neural and behavioural oscillations are influenced by respiratory state as breathing cycles from inspiration to expiration. To explore the mechanisms behind these effects, we carried out a psychophysical experiment where 41 participants categorised dot motion and facial emotion stimuli in a standardised discrimination task. When comparing behaviour across respiratory states, we found that inspiration accelerated responses in both domains. We applied a hierarchical evidence accumulation model to determine which aspects of the latent decision process best explained this acceleration. Computational modelling showed that inspiration reduced evidential decision boundaries, such that participants prioritised speed over accuracy in the motion task. In contrast, inspiration shifted the starting point of affective evidence accumulation, inducing a bias towards categorising facial expressions as more positive. These findings provide a novel computational account of how respiratory rhythms modulate distinct aspects of perceptual and affective decision-dynamics.

## Introduction

The interoceptive rhythms of the body are essential for maintaining homeostasis (Allen, 2020; Craig, 2003). While our breathing, heart-beat, and other visceral oscillations are critical for keeping us alive, less is known about how these rhythms influence our perception of the world. Indeed, until recently, breathing in the brain was thought to be largely constrained to basic processes concerning only the maintenance of respiratory drive and gas exchange (Heck et al., 2017; Herrero et al., 2017). This idea has recently been challenged by novel neurophysiological and behavioural discoveries (for reviews see, Allen et al., 2022; Brændholt et al., 2023). These studies have shown that respiratory rhythms drive alterations in both neural oscillations and behaviour across numerous tasks and animal models.

The finding that perceptual decision-making depends on the rhythms of breathing has been shown to span multiple different cognitive and sensory modalities. One recent study found that across visual, auditory, and memory tasks, reaction times differed substantially if participants responded during inspiration as compared to expiration (Johannknecht & Kayser, 2022). Other studies have found that inspiration increases the speed of emotional processing (Zelano et al., 2016), and that participants make more volitional movements during expiration (Park et al., 2020). These and many more recent reports suggest that respiratory rhythms play a fundamental role in shaping exteroceptive, affective (Bagur et al., 2021; Mizuhara & Nittono, 2023; Zelano et al., 2016), and cognitive behaviour (for reviews, see Allen et al., 2022; Azzalini et al., 2019; Brændholt et al., 2023; Heck et al., 2022).

Theoretical proposals suggest that this modulation operates by adaptive gain control through the respiratory tuning of neural excitability, a mechanism that is critical for evidence accumulation during perceptual decision-making (Allen et al., 2022; Aston-Jones & Cohen, 2005; Cheadle et al., 2014; Corcoran et al., 2018; Kluger et al., 2021). While some studies have demonstrated psychological and neural effects of the respiratory cycle that are broadly consistent with shifts in neural gain (Kluger et al., 2021, 2023), so far no study has directly examined the underlying cognitive mechanisms using computational modelling. As studies have highlighted consistent modulation of reaction time by respiratory rhythms (Johannknecht & Kayser, 2022), we hypothesised that breath-brain coupling may alter the underlying latent variables governing perceptual and affective decision-making.

To test this hypothesis, we conducted a psychophysical experiment in which participants performed stimulus discrimination tasks in the modalities of visual random dot motion and affective face perception. By modelling the influence of the inspiratory-expiratory cycle on perceptual choices with a hierarchical drift diffusion approach (Wiecki et al., 2013), we evaluated which specific decision variables in each modality were modulated by the breathing state. This enabled us to test whether respiratory state effects are primarily elicited by changes in stimulus processing or response execution, and to evaluate the cognitive mechanisms underlying these putative effects. We confirmed that participants respond more quickly in both domains when responding during inspiration, and also found that this acceleration arises from the modulation of distinct decision variables across these modalities. Our findings highlight unique mechanisms by which the respiratory cycle modulates visual motion and affective face perception.

## Results

We tested the effects of respiratory state on visual and affective processing using a perceptual decision making task which alternated between each stimulus modality across blocks. To maximise the ability to detect respiratory state effects on perceptual behaviour, all stimuli were individually thresholded using a Bayesian psychophysical approach (Kontsevich & Tyler, 1999). Random dot motion (RDM) stimuli were calibrated to the individual motion coherence threshold for discriminating upwards vs downwards motion (mean coherence threshold = 0.21, SD = 0.14). For the affective domain, we adapted a face affect discrimination (FAD) procedure (Wang & Adolphs, 2017), which utilised face morphing to present stimuli conveying a range of emotional expressions between 100% happy and 0% angry and vice versa. This enabled us to titrate stimuli to the individual threshold for perceiving a facial expression as happy vs angry (mean face morph threshold = 53% happy, SD = 9). Following this staircasing procedure, participants completed a one-alternative forced choice paradigm during presentation of 320 near-threshold stimuli in each modality.

We first assessed respiratory effects on choice behaviour. As the perceptual decision process unfolds over time, it is inevitable that different parts of this process will sometimes pertain to different respiratory states. To establish whether respiratory-behavioural coupling primarily depends on aligning perceptual or motor states with the respiratory cycle, we analysed trials independently grouped by respiratory state at stimulus presentation and at the time of response. In the visual motion modality, we found a significantly higher hit rate (HR) for responses made during expiration compared to inspiration (mean difference = 2.0%, 95% Confidence Interval (CI) = [4-0%], *p* = 0.044). This effect was not observed when grouping trials based on respiratory state at stimulus presentation (Bayes factor in favour of the null, BF01: 4.67). For FAD the proportion of happy vs. angry responses did not differ significantly for either stimulus or response grouping (BF01: 3.44 and 5.46 respectively). See ***Table S1*** for all statistics.

We next determined whether response speed for the two domains differed as a function of respiratory state, again using both stimulus and response based grouping of trials. Here we found a predominant effect for response-grouped median reaction times, such that responses were faster when they occurred during inspiration vs expiration for both RDM, mean difference = 18.9 milliseconds (ms), 95% CI = [30-10 ms], *p* < .001, and for FAD, mean difference 21.2 = ms, 95% CI = [30-10 ms], *p* = .001 (see Fig 2). In contrast, no effect for either modality was observed when analysing reaction times grouped by stimulus onset (BF01, RDM: 5.21, FAD: 3.4).

**Fig 1:**
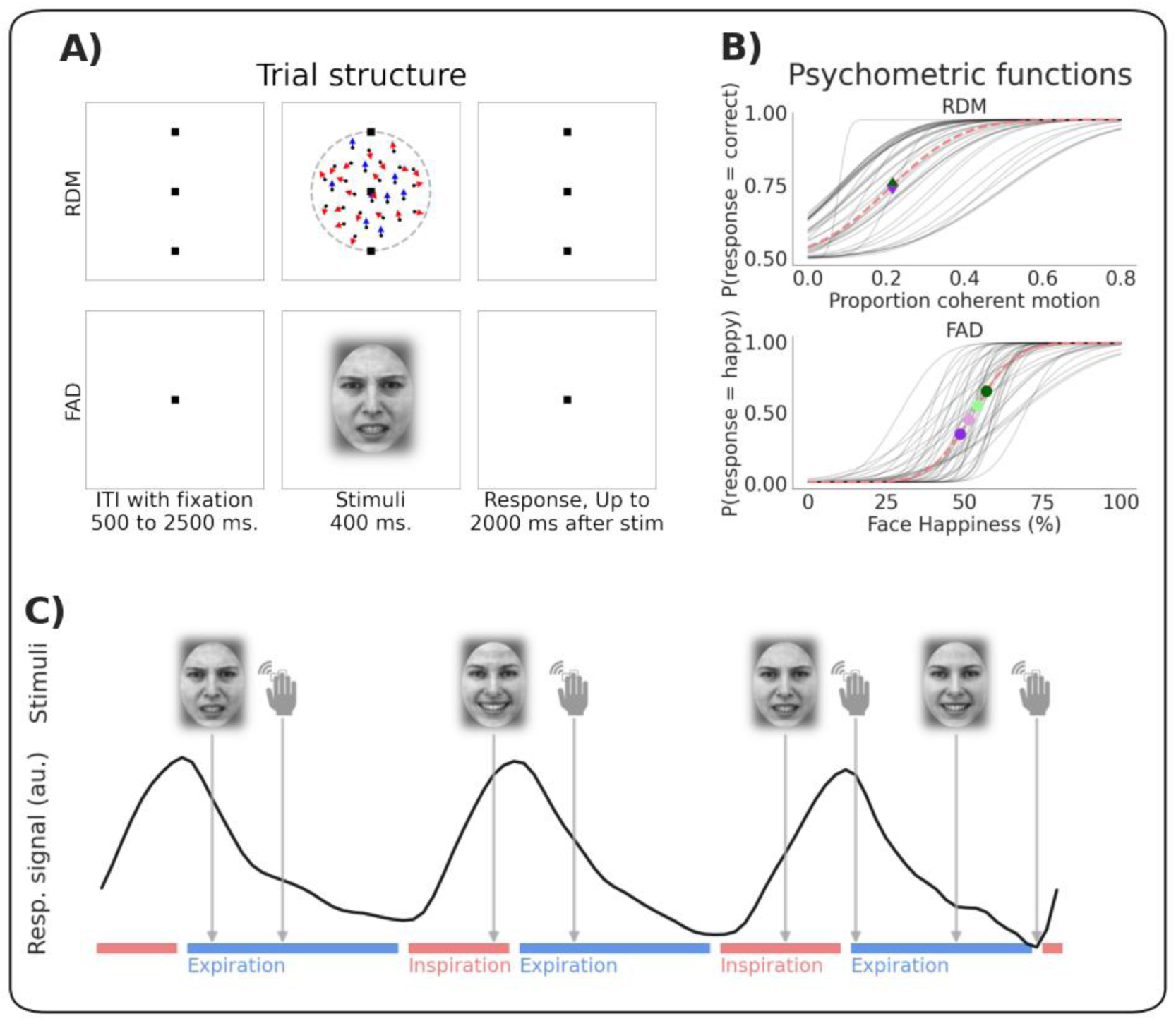
A) Trial structure of the random dot motion (RDM) and face affect discrimination (FAD) stimuli. B) Estimated psychometric functions for both RDM and FAD stimuli. Grey lines show individual participants’ traces, dashed red shows the group mean. Green and purple symbols indicate examples of the selected stimulus intensities used in the test trials. C) Each test trial was labelled both by the respiratory state at stimulus onset and at the time of response as indicated by the face and hand icons respectively. Face icons show examples of angry and happy stimuli. Red and blue lines represent periods of inspiratory and expiratory states respectively. ITI: Intertrial interval

**Fig 2:**
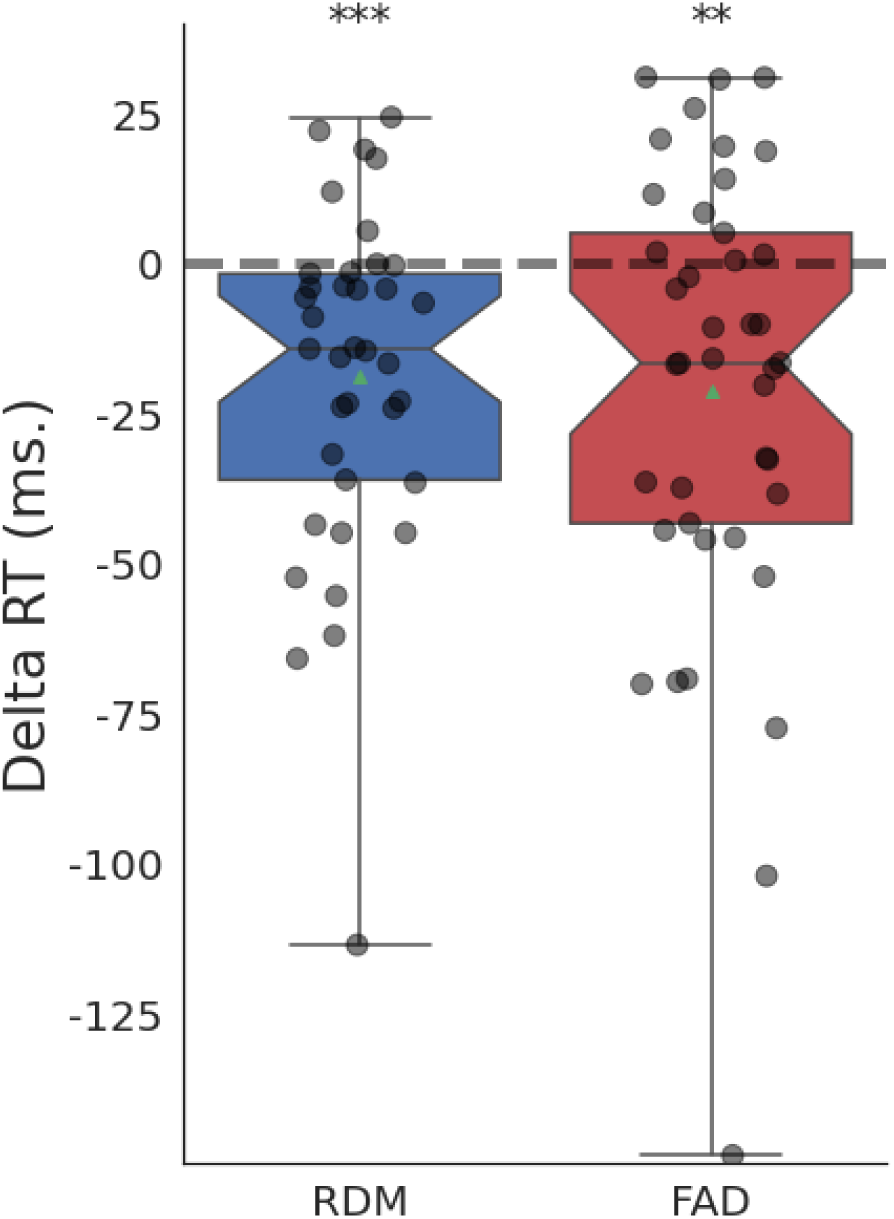
Boxplots depicting the difference (delta) in median reaction time during inspiration vs. expiration, based on response grouping for each domain. Positive values indicate higher values during inspiration compared to expiration. Dots indicate individual participant delta-medians. Notches indicate 95% CI of the median. RDM: Random dot motion, FAD: Face affect discrimination. **p < 0.01, ***p < 0.001

In summary, responses made during inspiration were faster for both modalities and less accurate only for the visual motion modality, pointing to a complex and heterogeneous coupling between respiratory state and decision-making. To establish the underlying mechanisms driving the observed coupling effects we employed drift diffusion modelling (DDM) (Ratcliff & McKoon, 2008). The drift diffusion model jointly analyses choice and reaction time data. The model postulates that speeded decisions can be characterised as a noisy accumulation of evidence originating from a starting point positioned between two decision boundaries (e.g., reporting a happy or an angry face), and additional latencies not related to the decision process itself. Fitting this model to choices, reaction times, and respiratory states allowed us to investigate whether the observed coupling effects were most effectively accounted for by changes in the rate of evidence accumulation (drift rate, *v*), participants’ biases towards particular responses (starting point bias, *z*), the balance between speed and accuracy (boundary separation, *a*), or by extra-decisional variables such as attention or motor preparation (non-decision time, *t*). For a visual representation of these parameters, see Fig 3A.

**Fig 3:**
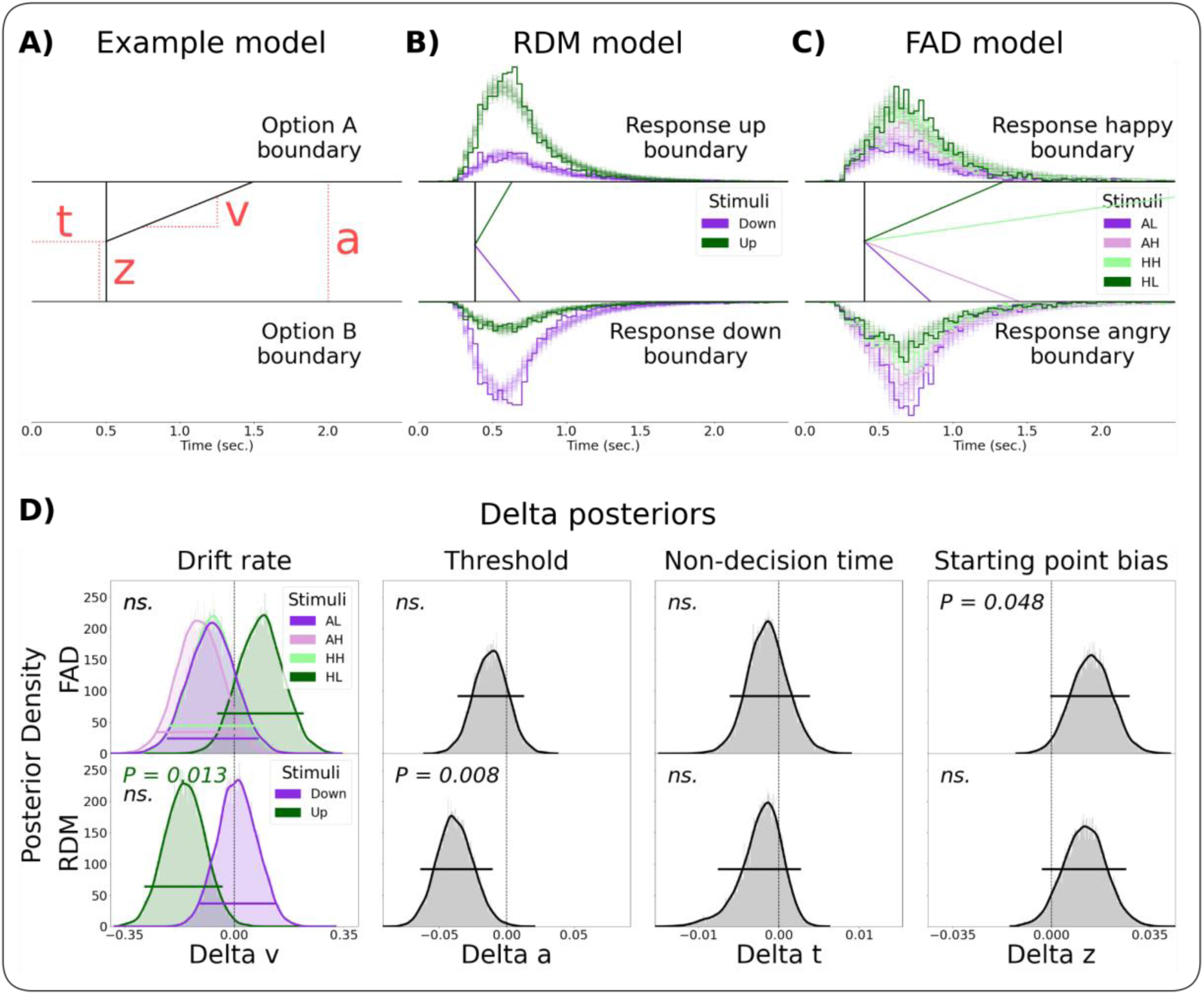
Drift Diffusion Modelling of Respiratory State Effects on Decision-Making. ***A)*** Schematic representation of the drift diffusion model (DDM) visualising the drift rate (v), non-decision time (t), bias (z), and decision threshold (a). ***B)*** Fitted group level DDM parameters and posterior predictive checks for the random dot motion (RDM) model and ***C)*** face affect discrimination (FAD) model. Drift rate was fitted for each stimulus class as shown by different coloured lines. Bold lined histograms represent the empirical reaction time histogram for each stimulus type and for each response option accumulated over time across all participants. Shaded lines represent the histograms of 100 data sets simulated based on the estimated model parameters. Note that the empirical and simulated data contains reaction times faster than the depicted group level non-decision time since some participants have shorter non-decision time than the group. ***D)*** Posterior predictive distributions for differences in DDM parameters by respiratory state. We found a significant reduction in decision threshold (*P* = 0.008) and drift rate of upwards stimuli (*P* = 0.013) for RDM, and a positive affective bias towards ‘happy’ responses during inspiration for FAD (*P* = 0.048). No significant shifts were found in drift rate of other stimuli, decision threshold in FAD or for non-decision time (all *P* > 0.05). Each parameter’s 95% highest posterior density interval (HPDI) is shown, reflecting the effect of respiration on decision dynamics. AL: angry, low ambiguity, AH: angry, high ambiguity, HH: happy, high ambiguity, HL: happy, low ambiguity.

We fit hierarchical DDMs where these parameters were modulated by the inspiratory vs respiratory state. Suitable model convergence and fit was assessed by calculating the Gelman-Rubin Statistic (see *Methods*), and by posterior predictive checks (**Fig 3B & C**). Inspection of the group-level posterior probability distribution (PPD) of each decision variable’s modulation by respiratory state (as illustrated in Fig 3D) revealed a significant decrease of boundary separation during inspiration for visual motion (*P* = 0.008), consistent with observed changes in speed and accuracy described above. For the affective modality, we observed a significant shift in starting point bias towards ‘happy’ responses during inspiration (*P* = 0.048). For both modalities, the state-related changes in evidence accumulation showed a complex, stimulus dependent pattern of respiratory modulation. That is, we found a significant reduction of drift rate for upwards motion stimuli (*P* = 0.013) during inspiration with no evidence for changes in drift rate for downwards motion stimuli (*P* = 0.889). For FAD we observed weak evidence that inspiration increased evidence accumulation for both low ambiguity stimuli (happy and angry) and for angry high ambiguity stimuli, as well as a weak trend towards a decrease in evidence accumulation for happy high ambiguity stimuli, however, none of these effects passed standard evidentiary boundaries and should thus be interpreted with caution.

These effects are illustrated in Fig 3D. Additionally, we report 95% highest posterior density intervals (HPDIs) and P-values in Table 1.

**Table 1:**
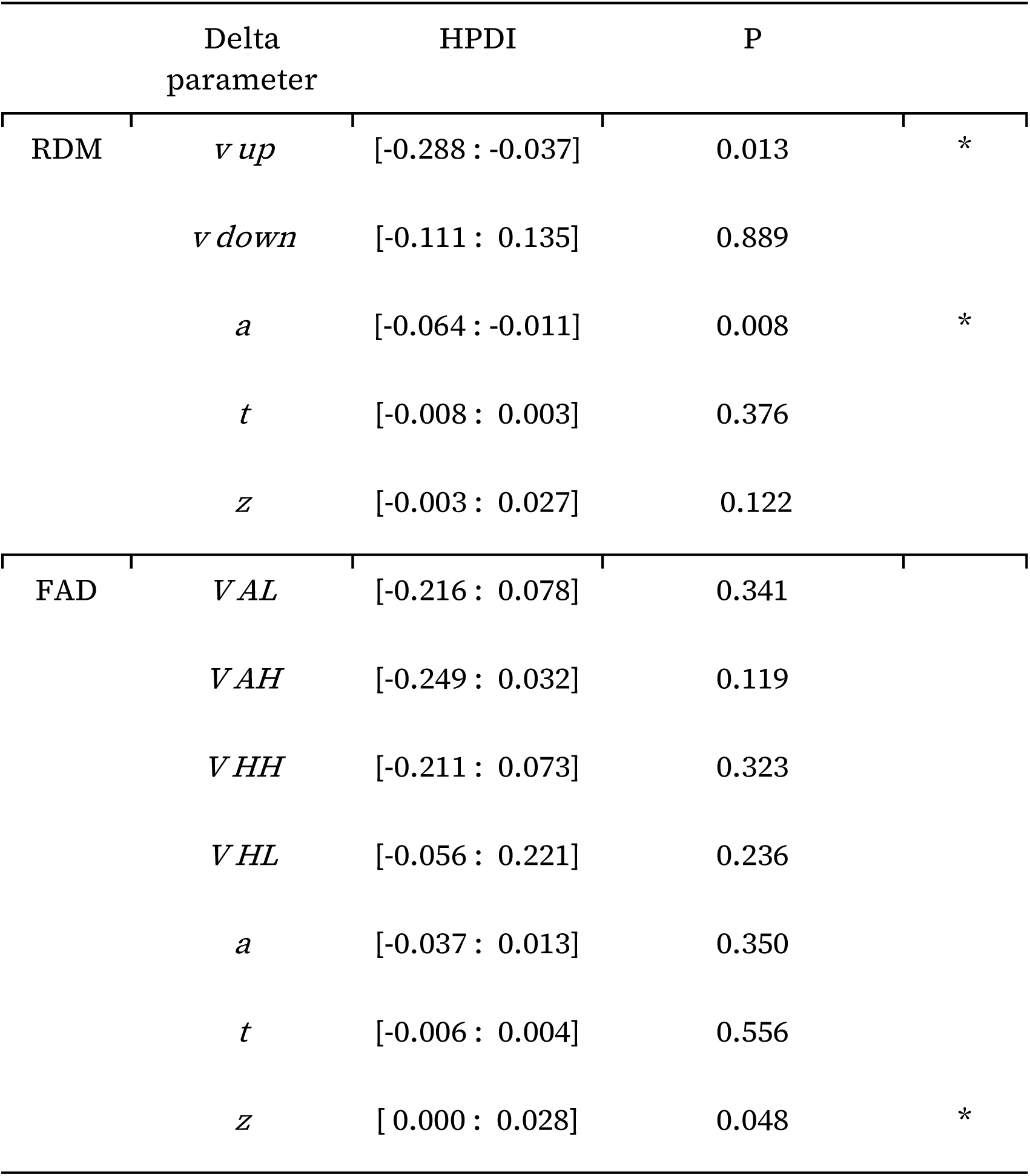
Highest posterior density interval (HPDI) and two sided Bayesian P-values (P) for the estimated respiratory state effects on decision making parameters. RDM: Random dot motion, FAD: Face Affect Discrimination, AL:angry, low ambiguity, AH: angry, high ambiguity, HH: happy, high ambiguity, HL: happy, low ambiguity. *P < 0.05

## Discussion

Numerous studies have demonstrated a close linkage between the rhythms which govern brain, body, and behaviour (Allen et al., 2022; Kluger et al., 2021, 2023; Varga & Heck, 2017). While the influence of respiratory states on cognition is by now well established, so far little direct evidence has emerged to explain the computational mechanisms underlying these effects. Here, we confirmed previous findings that inspiration during responses is associated with faster reaction times across visual and affective stimulus processing domains (Johannknecht & Kayser, 2022). However, through computational modelling, we found that these surface level phenomena are explained by distinct respiratory modulation of latent decision variables. Our study sheds new light on how respiratory-brain interactions control behaviour, and highlights the complexity of embodied modulation of exteroceptive perception.

In the context of visual motion discrimination, we found an inspiratory reduction of decision threshold. This indicates that responding during inspiration shifts decision-making to prioritise speed vs accuracy. Previous studies have indicated that respiratory state modulates cortical motor readiness (Park et al., 2020), cortico-spinal excitability (Li & Rymer, 2011) and cortico-muscular communication (Kluger & Gross, 2020), all of which point towards a possible effect of motor system activation. As such, one interpretation of our result is that the respiratory state can shift the threshold of executing actions, leading to shifts in both the frequency of spontaneous actions, as shown by Park and colleagues (2020), and in the amount of evidence needed to trigger a decision response, as our findings show. This suggestion is in line with recent developments in the DDM literature that increasingly poses motor areas not as simple effectors, subject to the output of higher order decision variables, but rather as important integrators of evidence, gating the decision and thereby contributing to setting the evidence threshold (Rogge et al., 2022; Steinemann et al., 2018). On this basis, we suggest that respiration may alter motor readiness during visual processing.

In the context of discriminating between angry and happy faces, we found a unimodal increase in the bias towards positive emotional judgments during inspiration. This selective modulation of affective decision bias by inspiration adds further nuance to the findings of Zelano et al. (2016) and Mizuhara and Nittono (2023), that reported modulation of affective processing by respiratory phase. In rodents, the respiratory rhythm has also been linked to the maintenance of fear-conditioned responses, suggesting a significant role for breathing in emotional regulation (Bagur et al., 2021). Our findings highlight a unique mechanism through which respiration may bias evidence accumulation to bring about these and other changes in affective decision-making.

Numerous authors have hypothesised that breath-brain coupling effects may depend in part on the modulation of neural excitation and perceptual tuning (Allen et al., 2021; Kluger et al., 2021, 2023; Varga & Heck, 2017). On these grounds, one might have expected to observe robust modulation of evidence accumulation across stimulus modalities. In contrast, here we report a complex pattern of effects with respect to drift rate, such that evidence accumulation is selectively enhanced or suppressed in a stimulus-specific manner. For example, for visual motion discrimination, we found that inspiration is associated with a decrease in the rate of evidence accumulation only for upwards motion stimuli, whereas drift was unaffected by respiratory state for downwards stimuli. For the affective modality, no significant changes were seen for any of the stimulus types. However, we observed trends towards increased evidence accumulation during inspiration for happy low ambiguity (HL) stimuli, as well as weak decreases in evidence accumulation for all other stimulus types. While these results should be interpreted with caution, they may indicate the possibility of stimulus specific neural tuning by the respiratory rhythm across perceptual modalities.

What neural mechanism could explain our findings? Breathing influences the brain through various pathways (see Allen et al., 2022 for review). Relevant examples include the brainstem mediated modulation of tonic noradrenaline (de Gee et al., 2017; Yackle et al., 2017), olfactory bulb propagated phase amplitude coupling (Biskamp et al., 2017; Cavelli et al., 2020; Ito et al., 2014; Zelano et al., 2016), as well as respiratory modulation of affective and prefrontal circuits (Bijanki et al., 2014; Duerden et al., 2013; Khalsa et al., 2016; Pichon et al., 2012). Other possible complementary mechanisms include respiratory coupled fluctuations in the physicochemical properties of the central nervous system, such as cardiorespiratory modulation of baroreceptors (Hamill, 2023) and blood gas concentrations (Salvati et al., 2022; Xu et al., 2011). Given the complexity and heterogeneity of these pathways, future studies will need to directly investigate the neural mechanisms underlying respiratory modulation of decision dynamics. For example, one recent study found that the flexible modulation of decision bias depends upon alpha-band neuromodulation (Kloosterman et al., 2019). An intriguing avenue for future research could be to combine these approaches, for example by examining the neural correlates of respiratory shifts in affective bias, which could depend upon a similar mechanism.

Our study has a few key limitations that could be addressed in future investigations. Participants viewed stimuli which were presented at peri-threshold levels. While this manipulation ensured that respiratory effects could be observed while controlling for stimulus specific biases, it potentially could introduce confounds related to drifting participant performance over time due to cognitive fatigue. While inspection of performance across blocks of trials suggested that performance was relatively stable, future studies could utilise adaptive thresholding, which would offer the further benefit of enabling estimation of respiratory modulated psychophysics (Kluger et al., 2021).

Another important limitation is that our focus on visual perception modalities inherently restricts the generalizability of our findings. Future research could overcome these limitations by employing varied experimental protocols and including a broader range of sensory modalities to enhance the applicability of the findings across different cognitive domains. Finally, while we here focus on the commonly reported effects concerning the binary treatment of the respiratory cycle, future studies could make use of more complex methods examining the link between constant respiratory phase and behaviour.

Finally, although our results suggest that breathing exerts unique rhythms on visual and affective perception, it should be noted that comparable, non-significant effects (e.g., bias in for RDM and threshold for FAD) can be observed in Fig 3. While our study utilised a relatively large number of trials and participants for a psychophysical design, it is possible that these effects would have been significant with more data. It may be in the end that the overall magnitude but not quality of these effects is differentially modulated; future studies could benefit from using a confirmatory (e.g., using registered reports) design with enhanced within and between-subject power to conclusively decide this issue.

## Methods

### Participants

Participants were recruited from a local online participant database, Sona Systems. A total of 41 healthy human participants completed the study (mean age = 26, SD = 6, 27 female). Participants received financial compensation for their involvement in the study. The study was conducted in accordance with the Declaration of Helsinki and was approved by the Region Midtjylland Ethics Committee.

### Experimental Setup

Participants sat upright at a desktop computer with their chins and foreheads stabilised by a forehead and chin rest to reduce respiratory-related head movements. The headrest and table height were tailored for comfort, setting an approximate 40 cm distance between participants’ eyes and a 24-inch LED monitor with a 60 Hz refresh rate and 1920×1080 resolution. Participants used the up and down arrow keys on a mechanical keyboard to input responses.

### Physiological Recordings

During the experiment, participants wore a respiration belt (Vernier, Go Direct Respiration Belt, GDX-RB), connected to the stimulus computer via a USB cable. The belt was individually adjusted and positioned at the mid-thoracic point, aligned with the lower end of the sternum, and continuously measured the force applied to it at a sampling frequency of 10 Hz. Simultaneously, a time series of experimental triggers was collected in order to align the respiratory data with stimulus and response onset.

### Stimuli

Motion perception was tested using random dot motion stimuli. A central fixation point and two reference points at 3° over and under the fixation respectively were displayed throughout RDM trial blocks (point size = 0.1°). Motion stimuli consisted of 1000 black dots on a grey background moving within a circular aperture 6° in diameter. Each dot was 1 pixel in size, moved at a speed of 9°/s and had a lifespan of 5 frames after which they were replaced randomly within the aperture. *Signal* dots comprised the fraction of dots which displayed coherent motion upwards or downwards, whereas *noise* dots moved in a new random direction every time they were replaced (see Fig 1A for an example stimulus).

The emotional stimuli comprised 201 morphed faces that transitioned in discrete steps from fully happy to fully angry. These were generated based on the Karolinska Directed Emotional Faces (KDEF) dataset (Lundqvist, D., Flykt, A., & Öhman, A., 1998). To minimise face-specific biases, we created two average anchor faces for happy and angry expressions using WebMorph (DeBruine, 2018). Nine individual faces with the highest normed ratings for happiness or anger in the KDEF dataset were averaged for each anchor. These faces met additional criteria, such as showing teeth in both expressions, as teeth have been identified as significant valence cues (Calvo & Marrero, 2009; Kohler et al., 2004). We then generated 201 morph levels between these anchor images. The low-level properties (i.e., spatial frequency, luminance and contrast) of the images were equalised across all images using the SHINE toolbox (Willenbockel et al., 2010). Specifically, the Fourier amplitude spectra were matched and the mean luminance and contrast were normalised. Each image was further processed with a Gaussian kernel, causing the face to gradually blur into the background.

### Task

For RDM stimuli, participants decided whether each stimulus was predominately upwards or downwards, with the level of motion coherence titrated to each individual’s threshold. For FAD stimuli, participants viewed titrated face morphs and decided whether each stimulus was more happy or angry. These stimulus levels were calibrated to each individual’s subjective perception, comprising four stimulus levels with two just above (happy, high and low ambiguity) and two just below (angry, high and low ambiguity) their affective perceptual threshold. The structure of the task consisted of a self-paced introduction to each of the two stimulus modalities followed by a staircase procedure establishing thresholds for each modality (see *Staircasing procedure,* below). After the thresholding procedure, participants completed four blocks of 80 stimuli of each modality, giving a total of 320 trials per domain. In all blocks the stimulus types were equally frequent and appeared in a pseudo randomised order, such that for RDM every 10 trials contained 5 upwards and 5 downwards stimuli, while for FAD each 8 trials contained 2 stimuli of each emotion level. Self-paced breaks separated the blocks. The blocks alternated between the two domains in a counterbalanced order between participants.

For both domains, the stimuli were presented for 400 ms. Responses were recorded from the stimulus onset and up to two seconds after the offset, followed by a variable inter-trial interval (ITI). To avoid spontaneous alignment of participants’ respiratory cycle to the frequency of stimuli presentation the ITI was jittered according to a uniform distribution ranging from 500 to 2500 ms (see Fig 1A). The experiment and stimuli were implemented using PsychoPy (Peirce et al., 2019).

### Stimulus Threshold Estimation

To ascertain participants’ perceptual thresholds for both domain, we employed a Bayesian staircasing procedure known as the ‘Psi’ method (Kontsevich & Tyler, 1999). This technique adaptively estimates the individual psychometric function, which represents the probability function correlating stimulus intensity with a specific binary outcome (see Fig 1B). Each staircase comprised 50 trials for both tasks, and in the event of missed responses, the trials were repeated with the same stimulus intensity.

For the RDM stimuli the threshold was defined as the coherence level at which participants had a 75% probability of providing a correct response. Psi parameters were as follows: intensity range: 0-1, threshold range: 0-0.5, slope range: 0.01-0.2, intensity precision: 0.01, threshold precision: 0.01, slope precision: 0.1, guess rate: 0.05, step type: ‘lin’, expected minimum: 0.5. The mean estimated coherence threshold across participants was 0.21, SD = 0.14. We then used the estimated psychometric functions to present dot-motion stimuli calibrated to each participant threshold in the RDM test-trials, resulting in an observed mean hit rate of 74% across participants.

For the FAD stimuli the threshold was defined as the point of subjective equality (PSE) of the emotional stimuli, corresponding to a 50% probability of responding ‘happy.’ The Psi parameters were as follows: intensity range: 0-200, threshold range 0-200 (i.e., corresponding to 0-100% happy in 0.5% steps), slope range: 0.001-50, intensity precision: 1, threshold precision: 1, slope precision: 1, guess rate: 0.02, step type: ‘lin’, expected minimum: 0. The mean PSE across participants was 53% happy, SD = 9. To ensure that participants did not view the same face stimulus repeatedly on test-trials, and thereby introduce variability across trials, we applied the estimated psychometric functions to identify four emotion stimuli centred around each participant’s PSE, corresponding to a probability of categorising a stimulus as ‘happy’ at 35% (angry low ambiguity), 45% (angry, high ambiguity), 55% (happy, high ambiguity), and 65% (happy low ambiguity) respectively. These stimulus levels were then presented during the FAD test-trials. This manipulation successfully manipulated affective perception, resulting in mean proportions of ‘happy’ responses at 32%, 42%, 53%, and 60% across the four stimulus levels.

## Analysis

### Behavioural and Physiological Preprocessing

Missed trials and trials with a reaction time < 100 ms were excluded from the analysis. Further, participants with extremely biassed responses, indicated by an absolute criterion > 0.6 in the RDM task, were excluded from RDM-related analyses. Criterion was calculated by averaging the inverse of the cumulative normal distribution for the proportion of correct responses to upwards-motion stimuli and the proportion of incorrect responses to downwards-motion stimuli and multiplying the result by negative one (Stanislaw & Todorov, 1999).

The respiratory signal was low-pass filtered using a 5th order digital butterworth filter with a cut-off frequency of 1 Hz (using the python scipy.signal.butter function), then standardised as z-scores. After preprocessing, each respiratory trace was visually inspected and segments with movement artefacts annotated. To extract the respiratory state from the preprocessed respiratory signal, peaks were identified using a peak detection algorithm (scipy.signal.find_peak in python with the parameters prominence: 0.2, distance: 10, width: 2). The minimum values between peaks were identified and labelled as troughs. See Supplementary online material for peak and trough detection and manual annotation for each participant.

The entire respiratory timeseries was then binarized into inspiratory segments, from trough to peak, and expiratory segments, from peak to trough. Stimulus onsets and responses which occurred exactly at the peak or trough were defined as relating to respiratory transitions and were not included in the respiratory state analyses, as these have been shown to represent distinct cognitive (Nakamura et al., 2018) and excitatory phases (Kluger et al., 2023), and there were too few in total (approx. 5% of total trials) to justify separate transition-based analyses (see Fig 1C).

Trials were then analysed separately grouped by respiratory state at stimulus onset and response, respectively. After behavioural and respiratory preprocessing a total of 37 RDM and 41 FAD datasets were available for analysis. For the included participants, a mean fraction of 7.6 %, SD = 2.6, of trials were excluded for RDM in stimulus grouping and 7.4 %, SD = 3.0, for response grouping. For FAD 7.9 %, SD = 3.3, of trials were excluded in the onset based analysis and 8.0 %, SD = 3.5, in the response based analysis. See Supplement table 2 for details on the exclusion steps.

### Paired T-tests

For RDM the hit rate (proportion correct responses), and for FAD the proportion of ‘happy’ responses, were calculated for each participant in each respiratory state for each of the two trial groupings. Similarly, the median RT was extracted for each of these trial groupings.

To test for an effect of the respiratory state on perceptual behaviour we employed paired t-tests for each of the outcome measures, comparing inspiration to expiration for stimulus and response grouping respectively. To evaluate the strength of evidence for any null effects, null Bayes Factors were calculated for each statistical comparison, using the default priors in the Pingouin analysis package.

All statistical analyses were conducted in Python using the Pingouin package v. 0.3.12 (Vallat, 2018).

### Computational Modeling of Respiratory State Effects

We used the HDDM python package, v. 0.9.7 (Wiecki et al., 2013), to fit hierarchical Bayesian DDMs. In the Bayesian framework, parameters are estimated as posterior probability distributions given the data and the priors. As no DDM of respiratory effects on perceptual behaviour has been performed so far, we used an uninformative, flat prior distribution for all parameters. The hierarchical nature of the model means that parameters are estimated both at group and participant-level. More precise participants have a greater impact on the group estimate, while simultaneously, the group estimate influences individual participant estimates, pulling them closer to the group mean. This partial pooling balances between and within-participant random effects, which improves both group (Wiecki et al., 2013) and participant-level parameter estimation (Katahira, 2016).

### Model Formulation

We fit choice and RT data from the two stimulus domains independently in a stimulus-coded response model, where drift rate is estimated for each stimulus class and where the response boundaries represent the two possible response categories (“upwards” vs. “downwards” and “happy” vs. “angry” respectively). For respiratory effects we added group level parameters signifying the difference in a given parameter during inspiration relative to expiration for all model parameters (*v, t, a* and *z*).

### Model Sampling

The models were estimated using Markov chain Monte Carlo sampling as implemented in the HDDM toolbox. Each model was sampled using 4 independent chains with 10.000 samples, a burn in of 2000 samples and a thin of 2. Convergence was tested by calculating the Gelman-Rubin statistics (Gelman & Rubin, 1992) (GRS), comparing within chain variance to between chain variance on all parameters and confirming that the GRS was less than 1.02 (Wiecki et al., 2013) for all parameters of all models and by visually inspecting the traces of all group level parameters (see *Supplement Fig 2*).

Further posterior predictive checks, plotting the actual RT distribution against simulated data based on the estimated parameters, were performed (see Fig 3B,C) to ensure the model captured key elements of the observed RT data.

### Parameter Tests for Effect of Respiratory State

We conducted Bayesian hypothesis testing (Cavanagh et al., 2014; Herz et al., 2016; O’Callaghan et al., 2017, 2021; Vandekerckhove et al., 2011) to assess the significance of respiratory effect parameters of the models, which represent differences from inspiration to expiration. For the PPDs of these parameters we calculated the probability mass suggesting a positive and negative effect respectively. Significance was determined based on whether two times the smallest of these probability masses (P) fell below 5%, corresponding to a two-sided test with alpha level = 0.05 (Murphy et al., 2014) (see Fig 3D).

