## Supplementary for "The respiratory cycle modulates distinct dynamics of affective and perceptual decision-making"

### Fig S1

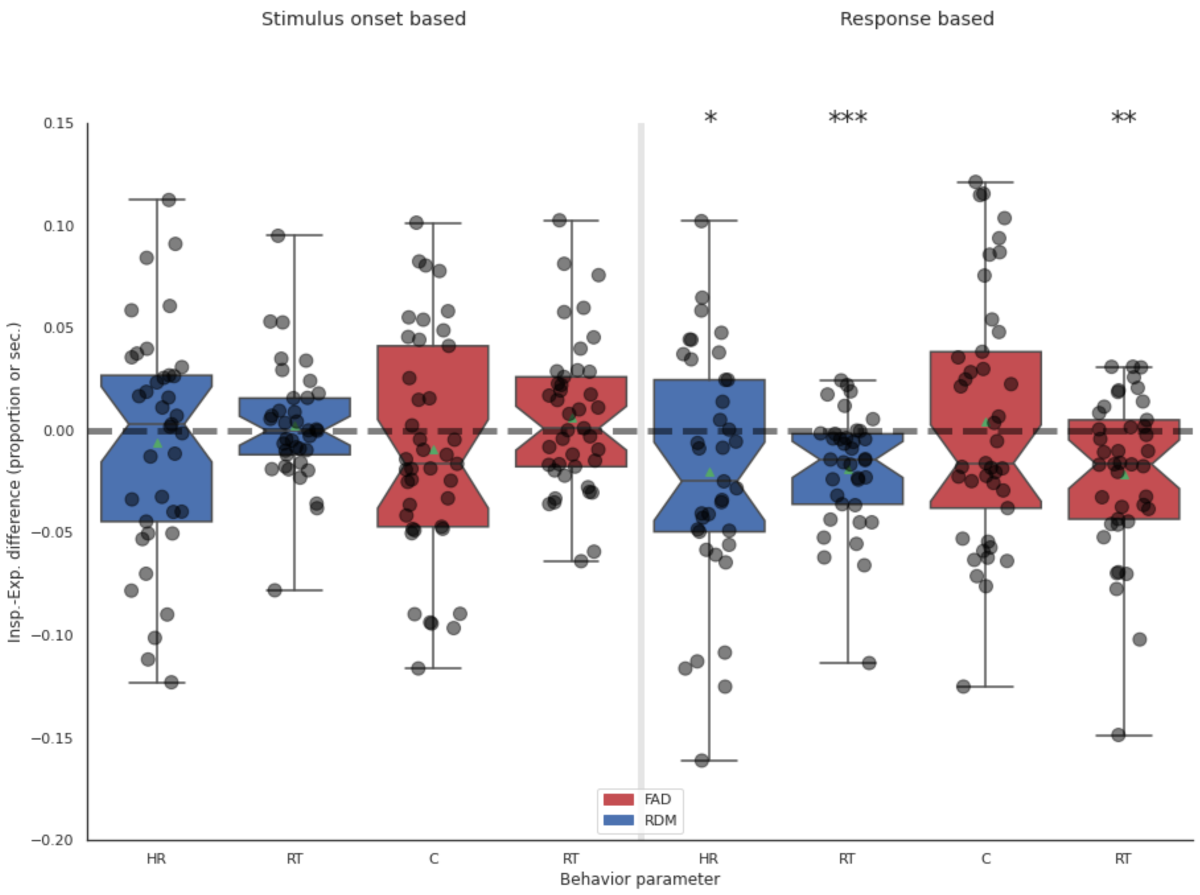


**Fig S1:** Difference in task behaviour during inspiration vs. expiration per participant. Positive values indicate higher values during inspiration compared to expiration. HR: Hit rate, RT: Reaction time, C: choice (proportion 'happy'-responses). Y-axis show difference in proportions for HR and C and difference in median RT for RT. Notches indicate 95%-CI of the median. RDM: Random dot motion, FAD: Face Affect Discrimination. *p< 0.05, **p< 0.01, ***p< 0.001

**Fig S2**

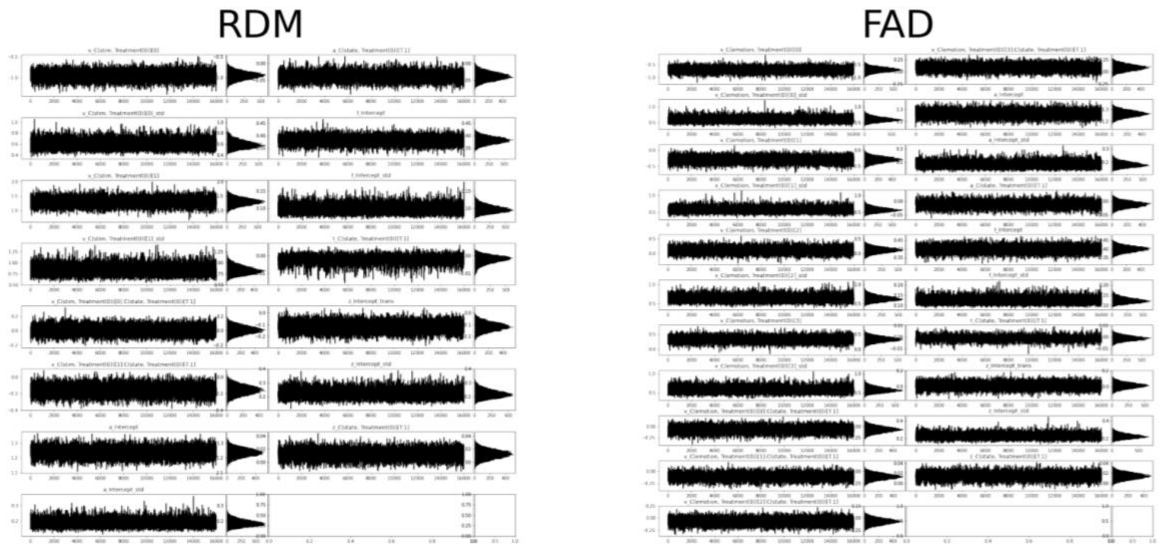


**Fig S2:** Caterpillar plots showing sampling traces for all fitted models and tasks. The plots show good model convergence across all chains.

**Fig S3**

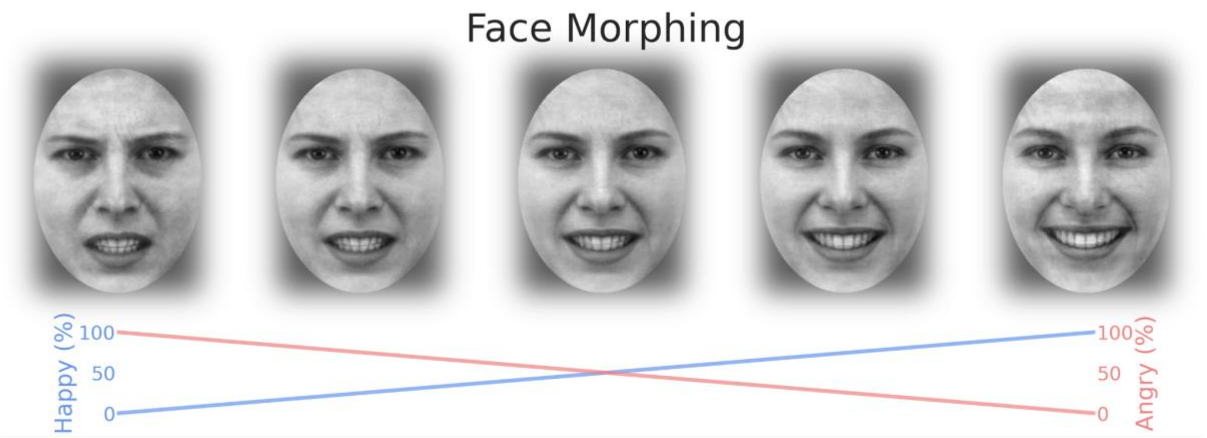


**Supplementary Fig 3:** Angry and happy face stimuli were morphed to create 201 categorical stimulus levels ranging from 100 % angry and 0 % happy to 100 % happy and 0 % angry.

### *Table S1: T-stats responses*

| Grouping | Task | Behav. | Diff. | 95%-CI | T | p-val. | BF10 | BF01 |
|---|---|---|---|---|---|---|---|---|
| Onset | RDM | HR | -0.0059 | [-0.02-0.01] | -0.6379 | 0.5276 | 0.2 | 4.673 |
| | RDM | RT | 0.0021 | [-0.01-0.01] | 0.4262 | 0.6725 | 0.2 | 5.208 |
| | FAD | C | -0.0093 | [-0.03-0.01] | -1.0815 | 0.286 | 0.3 | 3.436 |
| | FAD | RT | 0.0062 | [-0.01-0.02] | 1.0926 | 0.2811 | 0.3 | 3.401 |
| Response | RDM | HR | -0.0202 | -0.04--0.0] | -2.0885 | 0.0439 | 1.2 | 0.816 |
| | RDM | RT | -0.0189 | [-0.03--0.01] | -4.0651 | 0.0002 | 108.4 | 0.009 |
| | FAD | C | 0.004 | [-0.02-0.02] | 0.4184 | 0.6779 | 0.2 | 5.464 |
| | FAD | RT | -0.0212 | [-0.03--0.01] | -3.533 | 0.0011 | 28.9 | 0.035 |

**Table S1**: Statistical analysis of behaviour during inspiration vs. expiration. Diff. and 95%-CI refers to the mean difference between inspiration and expiration measured in proportions for HR and C and in seconds for RT. Positive values indicate higher values during inspiration compared to expiration. T and p-val. refers to paired t-tests. BF10 and BF01 is the Bayes factor in favour of the alternative and the null hypothesis respectively. Behav.: Behavioural parameter. HR: Hit rate, RT: Reaction time, C: choice (proportion ‘happy’-responses).

***Table S2: Trial Exclusion***

| Exclusion | RDM | | FAD | |
|---|---|---|---|---|
| Behav. | 0.4, 0.6 | | 0.7, 0.8 | |
| | Onset locked | Response locked | Onset locked | Response locked |
| Manual bad resp. signal | 1.7, 2.6 | 1.7, 2.6 | 2.1, 3.0 | 2.1, 3.0 |
| Trial on peak/trough | 5.4, 1.3 | 5.3, 1.7 | 5.0, 1.5 | 5.3, 1.5 |
| Total | 7.6, 2.6 | 7.4, 3.0 | 7.9, 3.3 | 8.0, 3.5 |

**Table S2**: Trial exclusion. For each task and trial grouping mean and SD in percentage points of excluded trials per step relative to the presented 320. Behav.: Exclusion based on behaviour, resp.: respiratory.
